# Neural volatility in orbitofrontal cortex underlies affective processing during healthy cognitive aging

**DOI:** 10.1101/2024.04.22.590523

**Authors:** Gargi Majumdar, Fahd Yazin, Arpan Banerjee, Dipanjan Roy

## Abstract

Understanding the mechanisms behind the variability of neural signals holds the key to the characterization of developmental and lifespan aging trajectories. Here we propose that tracking temporally structured neural fluctuations or volatility in brain areas during naturalistic tasks provides a more salient characterization of ageing trajectories compared to measures based on mean and resting-state variability. Compared to other prefrontal regions, the orbitofrontal cortices (OFC) exhibit higher BOLD signal variability in aged individuals. Neural latent state analysis revealed that, in contrast to the stable representations in younger adults, the OFC in older adults exhibited more temporally distorted representations, mirroring their distinct affective experiences. Furthermore, lower OFC volatility during the movie was associated with participant’s bias toward positive responses in a separate emotional reactivity task. This suggests orbitofrontal neural volatility might be a general adaptive response to affective computations. To investigate this further, a Bayesian learning model of valence dynamics was employed, which revealed that older adults exhibit heightened uncertainty in neural representations while estimating affective states. Collectively, these results indicate that neural volatility identified through BOLD variability carries unique information about older adults’ affective experiences and how naturalistic neuroimaging can chart a way forward in understanding this better.

## Introduction

Neural responses emerging from interactions within a complex system are inherently volatile^1,2^. Often, volatility is confused with “noise”; however, lately it has been recognized as a crucial cog of efficient neural processing^2–4^. From a complex system’s perspective, volatility is important in diverse fields such as financial markets and traffic systems and supports adaptability and resilience. This might support compensatory mechanisms reported widely in healthy human ageing^5–7^. A growing body of research has shown that neural variability plays a key role in higher-order cognition^8,9^, allowing for flexible navigation of task-specific demands^8,10,11^. We hereby define neural volatility as neural variability measured (standard deviation of neural signal) over a fixed period of time (under task, watching movies etc.). This is interchangeable in situations where the period of interest is equivalent to total experiment duration. This is strongly supported by theoretical frameworks suggest that neural variability reflects a statistically optimal representation of task uncertainty^12,13^. This interpretation of neural volatility as a cognitive facilitator is leveraged in aging neuroscience to compare how cognitive demands affect this measure across the lifespan^14–16^. However, studies often show highly region-specific^17^ and task-specific results^9,18^, making a generalized directional claim between neural variability and experience tenuous. This highlights the need to contextualize neural variability within specific regions and under relevant theoretical models. While studies on resting-state neural variability offer insights on spontaneous fluctuations, the absence of an explicit task make it difficult to link it to experience and specific theoretical frameworks outlined above^4^. In contrast, naturalistic neuroimaging offers a compelling and ecologically valid framework to explore how volatility to real-world stimuli can be used as functional markers of aging. Measures of volatility indexing subjective affective states across healthy aging can generate targeted task-specific biomarkers of pathological aging^19^.

A richly dynamic and socially complex movie requires one to flexibly navigate an array of internal and external affective states. Orbitofrontal subregions in the prefrontal cortex have been implicated in a multitude of affect-related studies^20,21^ and have strong theoretical underpinnings to learning^22^. Specifically, its medial and lateral sectors are thought to play complementary roles in affective computations^20^. The medial orbitofrontal cortex (mOFC) or ventromedial PFC is a core region involved in learning schemas^23^, state estimation^24^ and fluctuates between affective states^25^. In contrast, the lateral orbitofrontal cortex (lOFC) is mostly externally focused, predicts state transitions^26^, stimulus-outcome associations^27^ and is involved in predictive tracking of affect^20^. In short, both these regions show strong ties to affective experience in naturalistic experiences. Moreover, theoretical frameworks suggest that these regions are involved in estimating state uncertainty and its dysfunction can have implications in affective disorders^28^. Thus, having highly volatile neural activity in these regions with age should then exhibit broader changes to affective experience (Figure. 1a). For example, the mean neural activity might be similar in both young and older adults for a brain region. However, the neural variability, a proxy measure for capturing the volatility of the same region, can show distinctive patterns between the two groups. Such variability-based regional change would likely be specific to the ‘task-evoked’ volatility rather than intrinsic, resting-state variability^4,9^. Linking this to theoretical models of optimal task variable representations would offer testable model predictions to compare with neural data.

**Figure 1:**
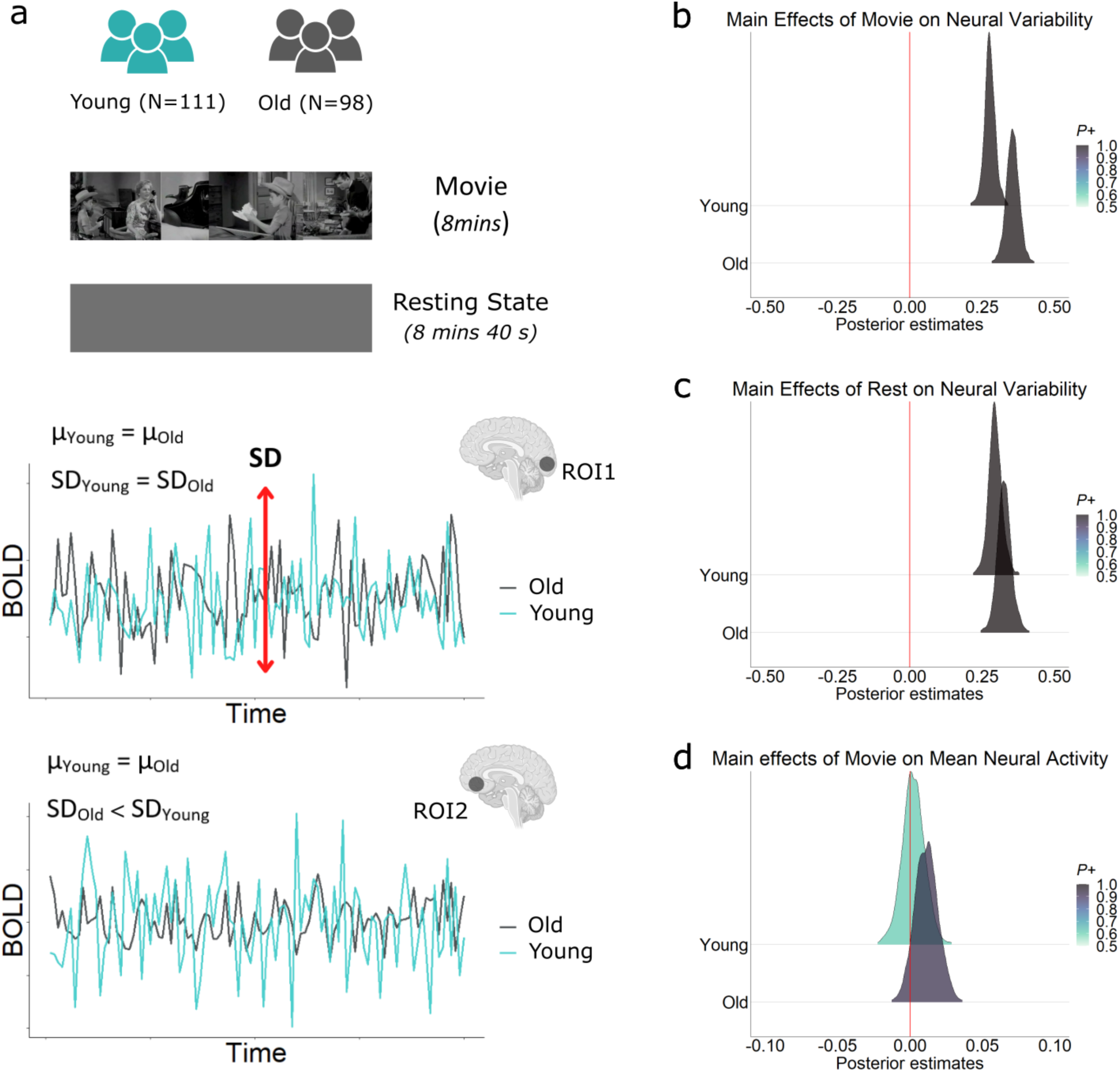
Neural volatility during naturalistic experience shows unique changes in aging. (a) Illustration depicting the regional differentiation based on neural variability across two age groups. We analysed average neural activity (µ_BOLD_) and the neural variability (SD_BOLD_) during movie-watching and resting state sessions in young (N=111, 18-35yrs) and older adults (N=98, 60-88 yrs). (Bottom) The illustration demonstrates that while the µ_BOLD_ and the SD_BOLD_ might be similar across age groups for ROI_1_, ROI_2_ shows differing levels of SD_BOLD_ between the age groups despite similar µ_BOLD_ values. This highlights that average neural activity and its variability provide distinct and complementary information into neural functioning. Main effect of (b) Movie and (c) Rest on SD_BOLD_ across young and older adults. Posterior density estimates show significantly more evidence for increased SD_BOLD_ in old than young (*P* = 1, BF = 415) in movie than that in the rest (*P* = 0.96, BF = 21.94) for the selected regions of interests. (d) Main effect of movie on µ_BOLD_. No significant difference was observed between the two groups (*P* = 0.83, BF = 4.89) for these regions.

In the present study, we explore how orbitofrontal neural variability during naturalistic viewing changes in older adults and contains unique information about computations underlying internal affective states. Using Bayesian modelling on large-scale neuroimaging data, we find that naturalistically evoked neural volatility showed a distinct increase in the cortex not observed in the mean BOLD responses or resting-state variability. Neural volatility was uniquely high in the OFC compared to other prefrontal subregions. Representations within the OFC were markedly distorted throughout the movie in the older adults, suggesting a divergence of affective experience relative to the young. We further link this neural volatility to the response bias of the participants towards positive affect. To assess whether this adaptation is suboptimal (computationally), we applied a Bayesian learning model, which revealed older adults incur more uncertainty in their estimates of emotional states. These findings, for the first time, link age-associated changes in neural volatility to shifts in subjective affective experience.

## Methods

### Participants

We divided our participant pool into two groups – Young (18-35 years) and Old (60-88 years). After preprocessing, a total of 209 individuals (Young: N = 111, µage = 28.5±4.9, Male = 48; Old: N = 98, µage = 77±5, Male = 45) were selected from the healthy population derived Cambridge Centre for Aging and Neuroscience (CamCan) cohort for this study^29,30^. We extracted the behavioral responses of these participants for the Facial Emotion Recognition (FER) and Emotional Reactivity and Regulation (EmoR) tasks. Owing to missing data, the final sample size for the FER was 203 (Y = 108, O = 95) and that of EmoR was 91 (Y = 49, µage = 28±5; O = 42, µage = 76.9±5.7). All participants provided their informed consent, and the data were collected under the Cambridgeshire Research Ethics Committee guidelines.

Additionally, we recruited independent groups of participants on an online platform (Prolific) to collect continuous ratings for arousal, valence and uncertainty for the same movie shown to the neuroimaging participants. 40 participants were recruited for each group (N=80), age and demographically matched with the fMRI individuals (UK). All participants were screened based on educational qualification and mental health status to reduce potential confounds and reported no previous history of watching this movie. The study was approved by the Institutional Human Ethics Committee of the National Brain Research Centre, India (NBRC).

### Stimulus

Our main behavioral and neuroimaging analyses were centered on a short, suspenseful movie called “Bang! You’re dead” (∼8mins) by Alfred Hitchcock.

### Continuous behavioral rating task

Four independent groups of participants continuously rated the movie for valence and arousal using a vertical slider on Prolific (Table 1). After rejecting subjects with missing data, this resulted in a final sample of 97 participants (Y: N = 34, µage = 24.5, Male =16; O: N = 30, µage = 67.8, Male = 14). For each group, we z-scored the participants’ responses and smoothed the averaged time courses using local regression smoothing (loess).

**Table 1.**
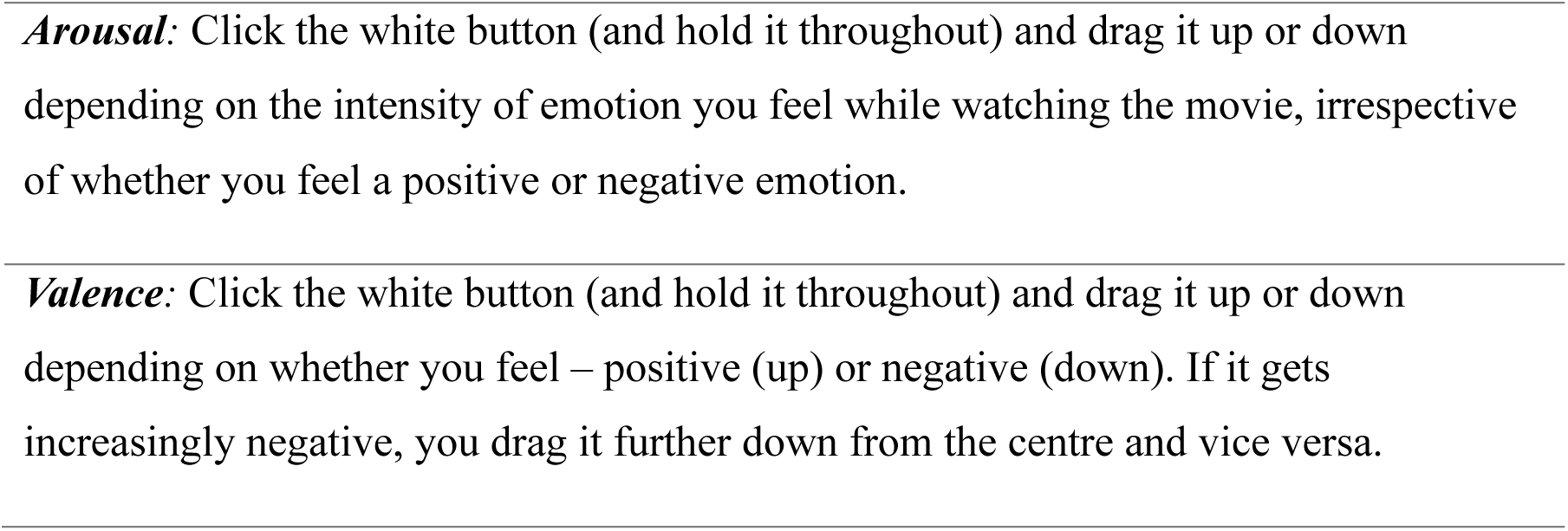
Instructions for the continuous rating task conducted in Prolific.

### fMRI acquisition and preprocessing

The fMRI data for the movie-watching and the resting state sessions were provided by the CamCan^29^. The functional scans for the movie-watching session were acquired using a T2∗-weighted echo planar imaging pulse sequence (TR = 2470 ms; TE = 9.4 ms, 21.2 ms,33 ms, 45 ms, 57 ms; 32 axial slices; slice thickness = 3.7 mm; FOV = 192 × 192 mm; voxel size = 3 × 3 × 4.44 mm), leading to a total of 193 volumes. The functional data for resting state was acquired using the same pulse sequence and acquisition parameters with a TR of 1970 ms. The acquisition time for this session was 8 min 40 s, which resulted in 261 functional volumes. The first 4 functional volumes of both sessions were discarded from the main analyses.

Neuroimaging data from both sessions were preprocessed in MATLAB (2019b) using Statistical Parametric Mapping (SPM12) (https://www.fil.ion.ucl.ac.uk/spm). The preprocessing steps included spatial realignment, slice-timing correction, co-registration of functional (T2) to anatomical (T1) scans, affine transformation to Montreal Neurological Institute template (MNI152), resampling to 2 mm isotropic voxels, and spatial smoothing with a 6 × 6 × 9-mm full-width at half-maximum Gaussian kernel. Whole-movie GLM was computed by regressing six motion parameters (X-, Y-, Z-displacement and pitch, roll, and yaw parameters) by least square regression. We excluded subjects with maximum framewise displacement >2.5 mm or 1.5° from further analysis. The extracted BOLD data were detrended and band-pass filtered (0.01–0.1 Hz) using a second-order Butterworth filter.

### Region-of-interest (ROI) definition

We focused our analyses on the prefrontal cortical regions due to their critical role in mediating higher-order cognitive functions, such as affect^20,31,32^, and their involvement in naturalistic tasks^33,34^.

For the neural variability analyses of the movie and resting state, we included 8 prefrontal ROIs – anterior cingulate cortex (ACC), dorsolateral PFC (dlPFC), dorsomedial (dmPFC), ventrolateral PFC (vlPFC), medial PFC (mPFC), medial orbitofrontal cortex (mOFC) and lateral orbitofrontal cortex (lOFC), posterior cingulate cortex (PCC). We also incorporated the visual cortex (VC) and Precuneus because of their roles in sensory and multi-sensory processing. All ROI masks were adapted from our previous work, where additional details can be found ^20^.

### Bayesian Hierarchical Regression (BHR) to model the effect of movie and resting state on neural variability and mean neural activity

First, we computed the neural variability as the standard deviation of the BOLD signal (SD_BOLD_) for the movie and the resting state. BOLD signals were extracted for each session from the residuals after regressing out signals from white matter, cerebrospinal fluid (CSF), and motion-related artifacts. Thereafter, the BOLD timeseries was normalized, filtered and centered for each ROI for both groups. We also followed an alternate pipeline to extract the BOLD which included applying ICA-AROMA before regressing out the CSF, WM, and Motion artifacts. Our region-specific result followed the same pattern, albeit with reduced differences (Extended data, Figure. 2-1). Furthermore, we also compared the ROI SD_BOLD_ values for both pipelines. We found similar patterns of increased variability for both the lOFC and mOFC, as well as an insignificant difference in VC (Extended data, Figure. 2-2). ICA-AROMA might be more conservative in removing movie-related signals, and thus, to preserve those for our further analysis, we instead followed our former pipeline, which follows as described in^19^.

**Figure 2:**
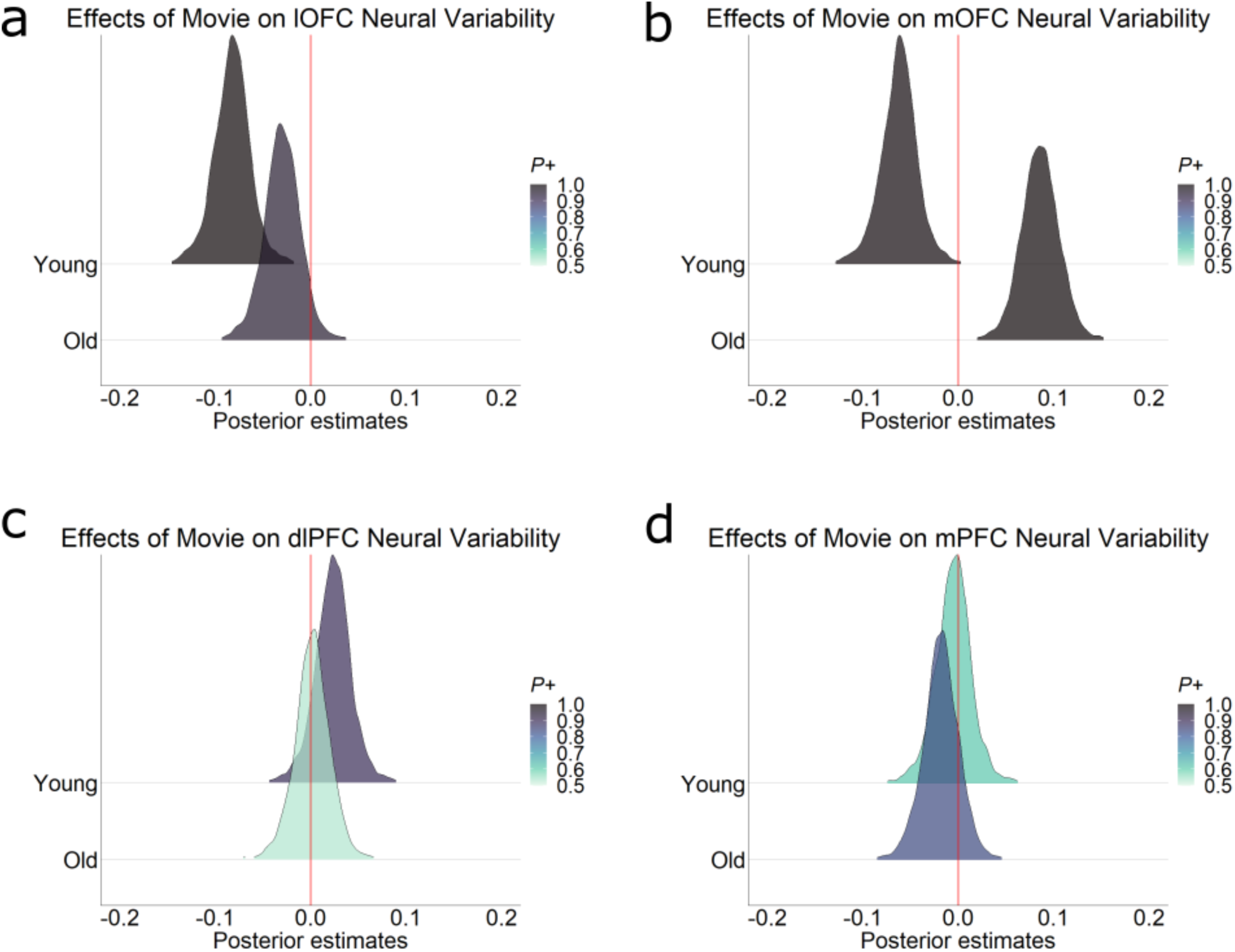
Increased neural variability shows in the Orbitofrontal cortex in aging. ROI-level effect of the movie on the SD_BOLD_ across the two groups. (a) Posterior density estimates of lOFC show significantly higher variability in old than young (Young: β_mean_ = −0.08, *P* = 0.99; Old: β_mean_ = - 0.03, *P* = 1). (b) Posterior density estimates of mOFC also shows increased variability in old compared to younger adults (Young: β_mean_ = −0.06, *P* = 0.99; Old: β_mean_ = 0.08, *P* = 0.99). No significant differences were observed in the neural variability between the two groups for (c) the dlPFC (Young: β_mean_ = 0.02, *P* = 0.89; Old: β_mean_ = 0.001, *P* = 0.53) and (d) mPFC (Young: β_mean_ = −0.005, *P* = 0.61; Old: β_mean_ = −0.18, *P* = 0.83).

Next, we modelled the effect of rest and movie with age on the BOLD variability of the participants, using a BHR (see Figure. 1b-d.). The dependent variable for the model input was given by the SD_BOLD_ of 10 ROIs, which included all prefrontal ROIs (mOFC, mPFC, dmPFC, lOFC, vlPFC, dlPFC, ACC, PCC) along with VC and Precuneus.

The SD_BOLD_ of each session (Rest/Movie) for each subject was modeled as a Student *t*-distribution: SD_BOLD_ = Student *t (v*, *μ*, *σ),*

where *v* is the degrees of freedom, *σ* is the scale parameter of the distribution, and *μ* is the mean which can be expressed as:

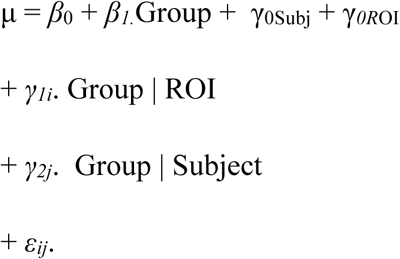

Here, *β*_0_ is the overall intercept, γ*_0R_*_OI_ is the varying intercept over *i* ROIs, γ_0Subj_ is the varying intercept over *j* participants. *β_1_* reflects the slope of the predictor, which is the group (Movie and Rest with Age; four groups). *γ_1j_* indicates the varying slope term for the group across each ROI. Likewise, *γ_2j_*, indicates the group slope varying across all *i* (=209) participants. *ε* connote the population-level variance terms. Weak, noninformative priors were applied over the model terms. This model incorporated the effect of rest and movie, while simultaneously accounting for within-group and between-group variability on the level of ROIs.

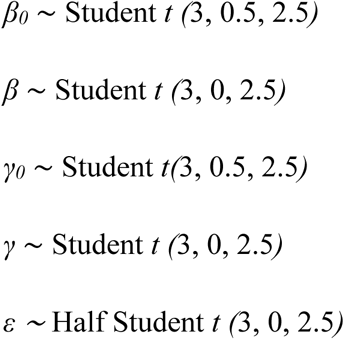

An evidence of *P >* 95% is equivalent to the frequentist depiction of significant *P*-values. In addition, we also quantified the effect size by computing the BF (calculated by the Savage-Dickey ratio).

A similar model was used to examine the effect of movie and resting-state conditions on mean neural activity across age groups, using the same ROIs. In this model, the standard deviation of BOLD (SD_BOLD_) was replaced with the average BOLD activity for each ROI.

### Temporal Potential of Heat-diffusion for Affinity-based Transition Embedding (PHATE)

Next, we sought to explore, whether the latent temporal trajectory in the OFC during movie-watching reflect the subjective emotional experience of young and older adults. For this, we implemented a non-linear, unsupervised dimensional-reduction technique called temporal PHATE (T-PHATE) ^35^. PHATE is recognized for its distinct advantages compared to conventional dimensional-reduction techniques like PCA and tSNE, as it effectively preserves both local and global data structures. Additionally, its non-linear nature and ability to capture the temporal evolution of a component present in the stimulus makes it well-suited for naturalistic paradigm.

It considers an input matrix of TR x Voxel, on which it computes the Euclidean distance D between two TRs, where D(*i*, *j*) = ||*X*_*i*_, − *X*_*j*_,||^2^. Then it converts this matrix into a local affinity matrix *K* using an adaptive bandwidth Gaussian kernel. The affinity matrix *K* is then row-normalized to get the initial probabilities *P*, which are used for the Markovian random-walk diffusion process. The diffusion operator *P* is then raised to the *t*_*D*_^th^ power, where *t*_*D*_ is the PHATE optimal diffusion time scale, to perform a *t*_*D*_-step random walk over the data. This specific diffusion process effectively infers more global relations beyond the local affinities. To further preserve both the local and manifold-intrinsic global structure of data, PHATE computes the diffusion potential distance *P*_D_ between the distribution at the *i*th row P*^tD^_i_* and the distribution at the *j*th row P*^tD^_j_*.

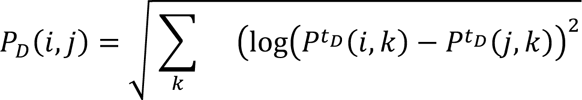

The potential distance is finally embedded with a distance embedding method (multidimensional scaling - MDS) for final visualisation. This embedding accurately visualizes trajectories and data distances, without requiring strict assumptions typically used by path-finding and tree-fitting algorithms. T-PHATE is a modified PHATE algorithm which is able to capture the temporal aspect of the data. The algorithm works the same as the original PHATE except for the temporal component infused in the diffusion matrix using the auto-covariance function in a Temporal Affinity Matrix^35^.

### Behavioral modelling - Hierarchical Gaussian Filter (HGF)

To investigate how the age-associated changes in task-evoked neural variability affect the computation during affective experience, we compared and contrasted multiple variants of the 3- level hierarchical Gaussian Filter (HGF)^36^. The HGF is a Bayesian learning model ^36^, which can be used to fit and recover parameters from a participant’s behavioral data or simulate models with different parameter configurations, scored on Log Model Evidence (LME). In brief, by employing variational Bayes, this model can infer an individual’s belief and uncertainty trajectory of hidden environmental states, which are modelled as a hierarchy of coupled Gaussian random walks. Since we did not have any behavioral response from the neuroimaging participants, we only used a perceptual model on a Bayes-optimal agent with the average valence ratings of each group as the input using the TAPAS toolbox (http://www.translationalneuromodeling.org/tapas/). With this input, the first layer, *x*_1_, models the probability of the valence at each time point. The next level, *x*_2_, estimates the underlying uncertainty associated with that of the predictions on the valence. Finally, the third level *x*_3_, estimates the overall volatility associated with the valence, i.e. the rate at which the belief on the uncertainty around valence changes. In this study, this level can signify the belief on the environmental stability or the context of the movie. These levels are related to each other with regard to the step-size of a Gaussian random walk. A prediction error i.e. a misestimation of the valence at the lower level will cause an update at the higher levels. These levels are coupled to each other by parameter ***κ***, which denotes a phasic, temporally varying parameter that determines the scaling of the volatility. Since we did not have any specific hypotheses on how this coupling might change with age, we kept the ***κ*** same for both groups. Our main hypotheses rested with the ***ω*** and ***ϑ*** parameters that denote a tonic, time-invariant parameter for the second and third levels, respectively. ***Ω*** captures the degree to which an agent updates his belief about the rate of change of valence 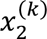 independent of the volatility and ***ϑ*** determines meta-volatility or the variance of the agent’s volatility.

The HGF is commonly implemented to dynamically model learning under uncertainty. However, previously this model had enabled us to uncover a predictive architecture where volatility trajectory estimated from the valence ratings reflected the subjective uncertainty of the participants^20^ during affective experiences. In this study, our goal was to test a few competing hypotheses, especially whether the older adults differed regarding 1) computation of the variance around the fluctuating valence across the movie and 2) updating their belief trajectory in the face of changing uncertainty on the valence. Thus, this can be construed as a form of model ‘lesioning’ wherein a normative model that explains a behavioral quantity and its latent parameters is altered, such that qualitative predictions similar to the data can be assessed.

To formalize these predictions, we simulated multiple models manipulating 2 free parameters ***ω*** and ***ϑ*** (Table 2), the tonic parameters for the second and third level, respectively. An increase in ***ω*** would indicate a higher uncertainty estimate (at the 2nd level) on the changing valence, whereas its decrease would indicate vice versa. The meta-volatility parameter ***ϑ***, influences the belief updates on the volatility (3rd level) of the valence. The resultant belief trajectories were visually inspected and after considering the Log-Model Evidence (LME), we selected the best model which could offer a parsimonious explanation of the age-associated changes during naturalistic experience in older adults.

**Table 2:** Prior parameters for all simulated HGF models.

| | $\omega_1 \mu$ | $\omega_2 \mu$ | $\omega_1 \sigma$ | $\omega_2 \sigma$ | $\theta \mu$ | $\theta \sigma$ | LME |
| --- | --- | --- | --- | --- | --- | --- | --- |
| Baseline Model | 99993 | -8 | 1 | 0.05 | 2 | 0.05 | 1703 |
| Increased Uncertainty at both levels | 99993 | -4 | 1 | 0.05 | 4 | 0.05 | 1717 |
| Decreased Uncertainty at both levels | 99993 | -10 | 1 | 0.05 | 0 | 0.05 | 1678 |
| Increased Uncertainty around Valence and decreased metavolatility | 99993 | -4 | 1 | 0.05 | 0.05 | 0.05 | 1737 |
| Decreased Uncertainty around Valence, increased metavolatility | 99993 | -10 | 1 | 0.05 | 4 | 0.05 | 1497 |

To analyse the effect of ***ω*** on the variance of belief around the changing valence we compared the belief update trajectories (ud2) of both groups.

### Hidden Markov Model (HMM)

Next, we wanted to explore how the age-associated computational changes are reflected in the latent state representations of the orbitofrontal cortex during naturalistic experience. For this, we fit an HMM to the individual subject’s BOLD time series of lOFC, mOFC, and VC (for control) for both groups (see Figure. 5e-f), generating a single emission sequence across all participants of a group. Multiple models were created from pre-specified states of range 2 to 10, which were then adjudicated by selecting the one with the least Bayesian information criteria (BIC). Parameter estimation for the models was done using the Baum-Welch algorithm (a variant of the Expectation-Maximization algorithm) by iteratively maximizing the expected joint log-likelihood of the parameters, given the observations and states. Decoding was done by the Viterbi algorithm by computing the maximum a posteriori state sequence for each of the HMMs. Thereafter, we compared the update trajectories from the HGF with the HMM state transitions of the orbitofrontal cortex subregions for the two groups.

### Behavioral tasks on Emotion processing

#### Emotional Reactivity and Regulation task

Our neural and behavioral data for the movie-watching task were derived from separate participant groups. We also wanted to examine whether the movie-driven neural variability reflect the affective profile of the same participants. For this, we used the data from an emotional reactivity task^29^ conducted with the same group of participants. Participants viewed short video clips of 30 seconds, chosen from a wide range of videos categorized into Positive (e.g. Babies playing), Negative (e.g. Accident) or Neutral (e.g. Workplace demonstration) clips. The presentation of the video was preceded by a cue to watch or regulate. For our study, we only selected the responses to the Watch condition, where participants were instructed to rate their emotional response on scales of 0 – 100 on how positive and negative they felt after each clip. Thereby, for each group, their behavioral ratings were classified into 6 types – Negative-watch-Negative (NegW_neg), Negative-watch-Positive (NegW_pos), Positive-watch-Negative (PosW_neg), Positive-watch-Positive (PosW_pos), Neutral-watch-Negative (NeuW_neg) and Neutral-watch-Positive (NeuW_pos). We compared the between-group variability of their responses for each condition.

To assess individual affective profile, we computed a positivity bias score. For affective clips, this was defined as the ratio of positive ratings to negative ratings across affective conditions— specifically, positive ratings for negatively cued clips (NegW_pos) divided by negative ratings for positively cued clips (PosW_neg). For neutral clips, the positivity bias was calculated as the ratio of positive to negative ratings (NeuW_pos / NeuW_neg). These two bias scores were analyzed separately and also averaged to create a combined positivity bias score. Finally, we correlated these behavioral positivity biases with movie-driven neural variability, focusing on the orbitofrontal cortex (OFC) as well as other prefrontal cortex (PFC) regions of interest (ROIs).

#### Facial Emotion Recognition task

Participants were presented with a series of emotional faces and for each trial, they had to characterize the emotion from six emotion categories – happy, sad, anger, fear, disgust, and surprise. To investigate potential misrepresentations of emotion, possibly stemming from increased valence variability, we generated co-occurrence matrices for the two groups across these 6 categories. Each point in the matrix represented the counts of responded emotions falling within the true emotion category, with diagonal elements set to 1 to signify instances where the responded emotion matched the true category. This was visualized as a chord diagram with a threshold of 1.6. Furthermore, we investigated the within-group and between-group behavioral variability (SD) across all categories and for each category, respectively.

## Results

### Age-associated changes in cortical neural volatility during naturalistic experience is distinct from resting-state variability

We aimed to investigate how the neural volatility alters with aging in response to a naturalistic movie (Figure. 1a). For this, we first modelled the BOLD by regressing out the averaged CSF and white matter timeseries, along with the 6 motion parameters. The residuals were used for further analyses^19^. We postulated that the neural volatility can be quantified using measures of neural variability as the standard deviation of the normalized, filtered, and mean-centered BOLD time series (SD_BOLD_) for both young and older adults for the movie (∼8mins) and the resting state (∼9mins) conditions. This enabled us to assess whether age-associated changes in neural variability reflect intrinsic traits or are induced by the affective states elicited by the movie.

We focused our analyses primarily on the prefrontal regions given their role in higher-order cognition and affective processing as well as evidence from our previous study^20^. The regions of interest included the prefrontal regions – anterior cingulate cortex (ACC), dorsomedial cortex (dmPFC), dorsolateral prefrontal cortex (dlPFC), medial prefrontal cortex (mPFC), ventrolateral prefrontal cortex (vlPFC), lOFC and mOFC, visual cortex (VC), Precuneus (Prec) and posterior cingulate cortex (PCC), the latter two being core Default Mode Network regions. Next, we constructed a Bayesian hierarchical regression with the SD_BOLD_ from the movie and the rest as the dependent variables. The model also included random effects of participants including random slopes for the fixed effects to capture participant-level variance, across movie and rest. This allowed us to assess whether the BOLD variability associated with these regions, is driven primarily by structural changes with age, in which case a similar pattern of change would be observed for both movie and resting-state conditions. Posterior evidence for each hypothesis was quantified through Bayes factors. We observed a significant increase in the SD_BOLD_ for the older individuals for the movie and not the rest (Movie: *P* = 1, BF = 415; Rest: *P* = 0.96, BF = 21) (Figure. 1b, c).

### Age-associated neural volatility offers unique information not captured by mean neural activity

One possibility is that the BOLD signal varies as a function of age affecting both the mean activity and variability of the young and older adults during movie-watching. This led us to investigate if the neural variability reflects a distinct source of information in contrast to the mean neural activity between the two groups. We computed the averaged BOLD signal for the movie-watching condition for the same ROIs and applied a Bayesian hierarchical regression using the previous model structure. No significant difference was found between the mean BOLD activities of the two groups (Figure. 1d) (Movie: *P* = 0.83, BF = 4.89).

The above results suggest two key points. First, the increase in cortical BOLD signal variability observed in healthy aging represents a movie-driven effect unobserved in the resting state. Second, within naturalistic settings, neural variability provides distinct insights than those captured by the mean neural activity.

### Increased movie-evoked neural volatility within lOFC and mOFC in older adults

Our findings so far suggest a distinct change in the movie-induced neural volatility between the two groups. Next, we wanted to focus on how these changes manifest at the regional levels and affect their corresponding functions in naturalistic experience. We particularly focused on the two subregions of the OFC known to mediate affective computations in naturalistic experiences^21,37,38^. This is also evidenced from our previous study which identified a distinct role of the lOFC and the mOFC in mediating affective experience during this movie^20^. We compared the SD_BOLD_ of the lOFC and mOFC between the young and older adults during movie-watching. We observed a similar pattern in the neural variability of these two regions with age (Figure. 2a,b). Older adults exhibited significantly higher SD_BOLD_ in the lOFC (Figure. 2a) (Young: β_mean_ = −0.08, 95% HDI = [-0.12, −0.04], *P* = 0.99; Old: β_mean_ = −0.03, 95% HDI = [-0.07, 0.007], *P* = 0.94), as well as the mOFC (Figure. 2b) (Young: β_mean_ = −0.06, 95% HDI = [-0.10, −0.02], *P* = 0.99; Old: β_mean_ = 0.08, 95% HDI = [0.04, 0.12], *P* = 0.99). Since OFC is particularly susceptible to head motion-related artifacts, we also investigated this following a more stringent preprocessing pipeline by applying ICA-AROMA before calculating BOLD variability. This was done to ensure that the observed differences were not mainly driven by the motion-related artifacts in older individuals (Extended data Figure. 2-1). Notably, we observed a similar pattern of increased neural variability for both the OFC subregions even after this correction.

Next, we sought to determine whether the movie-driven changes in neural variability of OFC observed in older adults reflected a general aging effect that could also be captured by the mean BOLD activity. For this, we compared their mean neural activity for the movie-watching condition for both lOFC and mOFC. No significant differences were found between the groups in the BOLD means for either the lOFC (Extended data, Figure. 1-1) (Young: β_mean_ = 0.003, 95% HDI = [-0.009, 0.02], *P* = 0.70; Old: β_mean_ = 0.002, 95% HDI = [-0.008, 0.01], *P* = 0.67) or mOFC (Young: β_mean_ = −0.004, 95% HDI = [-0.01, 0.009], *P* = 0.74; Old: β_mean_ = −0.005, 95% HDI = [-0.02, 0.006], *P* = 0.81).

Additionally, we also compared the SD_BOLD_ of dlPFC and mPFC between young and older adults, to assess the age-related changes in the neural variability of other prefrontal regions. These prefrontal regions are often involved in attentional control and social processing respectively, both of which can change with age, and thus relevant in this context. No significant differences were observed in the neural variability between the two groups for these regions (Figure. 2 c,d) (dlPFC: Young: β_mean_ = 0.02, 95% HDI = [-0.02, 0.06], *P* = 0.89; Old: β_mean_ = 0.001, 95% HDI = [-0.03, 0.03], *P* = 0.53; mPFC: Young: β_mean_ = −0.006, 95% HDI = [-0.05, 0.03], *P* = 0.61; Old: β_mean_ = - 0.02, 95% HDI = [-0.05, 0.02], *P* = 0.82). We observed a similar pattern of neural variability between the groups for these regions using the ICA-AROMA pipeline as well (Extended data, Figure. 2-1c, d).

The above results indicate that regional BOLD variability provides distinct neural signatures of aging that are not captured by mean BOLD activity alone. Secondly, the increased neural variability trends uniquely observed in the OFC suggests an age-specific shift in regional volatility during naturalistic stimulation, potentially reflecting altered affective processing in older adults.

### Disrupted temporal dynamics in OFC reflect volatile emotional experience in aging

Our findings so far suggest that older individuals exhibit an increased neural volatility in the lOFC and mOFC during naturalistic experience. But we have not yet directly examined if these regional changes can be linked to their affective experience. To explore this, we collected continuous ratings of Arousal for this movie from two independent groups of young and older adults, since the fMRI participants were freely viewing. Arousal here reflects the emotional intensity of the movie. We found a significant difference between their arousal ratings, with older adults displaying lower and more volatile arousal (Figure. 3a) (*r* = 0.67, *p* < 0.001). This effect was most pronounced during emotionally intense scenes, as rated by the young group.

**Figure 3:**
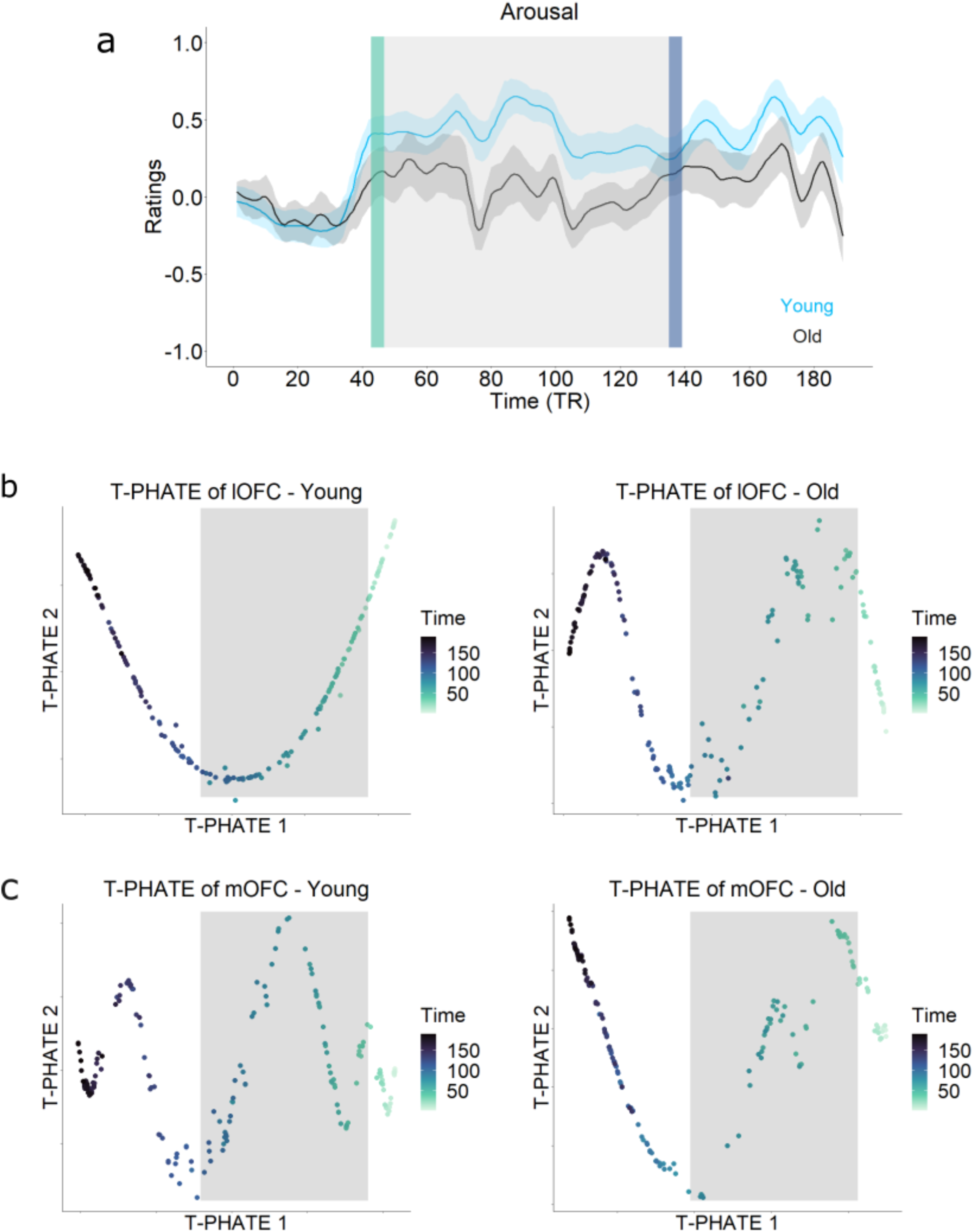
Changes in Orbitofrontal neural manifold trajectories reflect broad changes in affective experience. (a) Older adults exhibit lower and more variable arousal ratings than younger adults (r=0.67, p<0.001). Manifold embedding of the (b) lOFC and (c) mOFC using T-PHATE, with each data point indexed by TR. The latent temporal trajectory of both lOFC and mOFC show a conserved and sequential representation during a movie-watching task in young. In contrast, the older adults exhibit a disrupted latent trajectory, indicating alterations in their temporal dynamics in OFC, especially the lOFC. Notably, the disrupted segments in their latent trajectory correspond to their differential emotional experiences during those specific parts of the movie with lower and more variable (colored bars in the ratings correspond to the TRs in T-PHATE embeddings). Importantly, despite the neural and behavioral data being from two independent groups of participants, a similar pattern is observed in both.

The increased neural volatility exhibited in lOFC and mOFC may reflect deeper alterations in their temporal dynamics, particularly in naturalistic contexts like movie-viewing, where evolving experiences are shown to be encoded in their latent neural trajectories^39,40^. Changes to these latent temporal dynamics between the two groups could then be reflective of their experiences. To this end, we leveraged a non-linear dimensionality reduction method, Temporal-PHATE, on the BOLD timeseries of these regions (see Methods for full description). This method has been shown to capture the sequential progression of neural activity onto a low-dimensional space, revealing how these regions transition across time. This would ‘recover’ the temporal latent state trajectories that these regions would navigate during naturalistic experience on a group level.

In young adults, the latent representation in the lOFC (Figure. 3b left) and mOFC (Figure. 3c left) followed a preserved temporal sequence during movie-watching. Older adults, by contrast, revealed a temporally disrupted sequence (Figure. 3b, c, right). Crucially, the arousal responses showed the greatest differences between the groups at the time points where the latent sequences in the orbitofrontal cortex (OFC) were most disrupted in the older adults. This consistency between neural data and behavioral ratings, despite originating from two distinct groups, underscores the reliability of these results.

Overall, these results suggest that movie-evoked neural volatility in orbitofrontal region responses changes during aging could be a readout of the affective experience.

### Increased orbitofrontal neural volatility is associated with reduced positivity bias with age

The above findings suggest that orbitofrontal neural volatility in older adults during movie viewing may be linked to their emotional experiences. Since our neural and behavioral data were derived from separate participant groups, we next examined whether the movie-driven neural variability reflects the affective profile of the same participants. This is crucial to assess the generalizability of our findings. For this, we used the data from an emotional reactivity task^29^ conducted with the same group of participants. Briefly, participants watched short video clips (∼30s) of Neutral, Negative, and Positive valence and provided positive and negative ratings (0-100) of their emotional experience for each clip. We focused on the “Watch” condition to match the passive nature of movie-watching. First, we explored the behavioral variability in emotional reactivity between the two groups. We only focused on four conditions – Neutral watch positive (NeuW-Pos), Neutral watch negative (NeuW-Neg), Negative watch positive (NegW-Pos) and positive watch negative (PosW-Neg). Consistent with prior findings, all 4 conditions (see Methods for details) showed a consistent increase in variability in old irrespective of the valence (NegW_pos: SD_Y_ = 0.47, SD_O_ = 1.75; PosW_neg: SD_Y_ = 0.74, SDO = 1.47; NeuW_pos: SD_Y_ = 1.18, SDO = 2.29; NeuW_neg: SD_Y_ = 0.40, SD_O_ = 1.42) (Figure. 4a).

**Figure 4:**
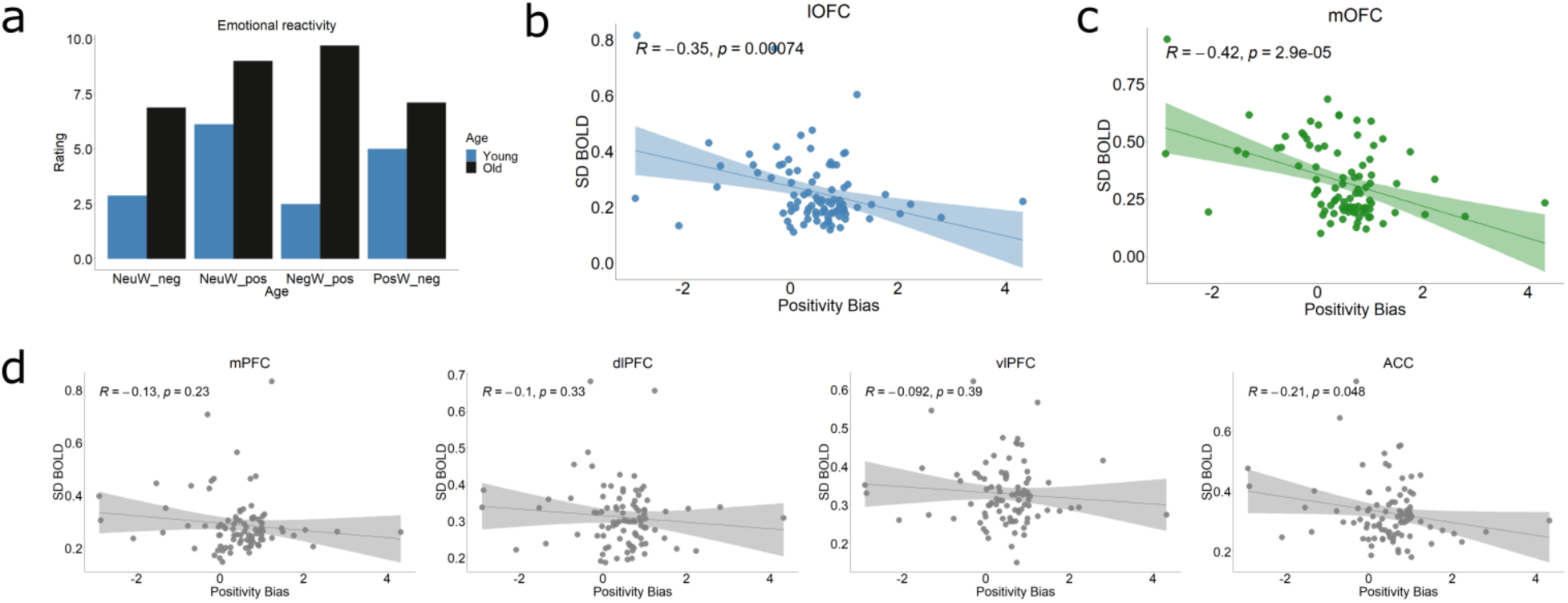
Increased neural variability in the orbitofrontal cortex is associated with reduced positivity bias in aging. (a) Between-group variability during the Emotion Reactivity task reveals greater SD in ratings among older adults compared to younger adults across all conditions. Participants viewed three categories of emotional clips (Neutral, Positive, and Negative) and rated them on both positive and negative affect scales, yielding six conditions in total. Four of these conditions (NeuW-Pos, NeuW-Neg, NegW-Pos, and PosW-Neg) were selected to compute a composite positivity bias score for each group. (b–c) Correlation plots showing that greater neural variability in the lOFC and mOFC is significantly associated with reduced positivity bias across age during the Emotion Reactivity task. (d) In contrast, other prefrontal regions (mPFC, dlPFC, vlPFC) commonly implicated in affective processing and regulation did not exhibit any such pattern in their SD_BOLD_ positivity bias. The SD_BOLD_ of ACC shows a marginally significant negative correlation with their positivity bias, though weaker than that observed for both lOFC and mOFC.

A substantial body of aging research suggests that older adults often exhibit a ‘positivity bias’ in emotional processing tasks ^41,42^, favoring positive over negative emotions, though this effect is not universally observed^43^. This led us to explore whether the neural variability in the OFC was associated with each participant’s bias to positive affect. We quantified each participant’s positivity bias as the ratio of their positive to negative ratings across neutral and affective clips (See Methods for details), with higher values indicating a greater bias to positive affect.

Across participants, we observed significant negative correlations between positivity bias and neural variability in both the lOFC (r = –0.35, p < 0.01) and mOFC (r = –0.42, p < 0.001) (Figure 4b, c). To assess the specificity of this relationship within the prefrontal cortex, we examined additional regions. No significant associations were found in the mPFC (r = –0.13, p = 0.23), dlPFC (r = –0.10, p = 0.33), or vlPFC (r = –0.09, p = 0.39), and only a marginal effect was observed in the ACC (r = –0.21, p = 0.048) (Figure. 4d). Importantly, this relationship between OFC neural variability and positivity bias remained significant when analyzing neutral and affective clips separately (Extended data, Figure. 4-1).

These results show that lower orbitofrontal neural volatility was linked to higher bias towards positive affect. This links neural variability in the OFC to an individual’s general affective computations, suggesting it might reflect an adaptive response of healthy aging.

### Older adults represent larger uncertainty during estimating emotional states

Increased neural variability has been suggested to facilitate dynamic behavioral switching based on task-specific demands^4,14,44^. Parallelly, theoretical frameworks propose that neural variability supports efficient encoding of stimulus-specific uncertainty during tasks^13,45^. Our previous study elaborated the role of orbitofrontal regions in encoding uncertainty during affective processing^20^. Specifically, we have shown that these regions are involved in optimally encoding the uncertainty around the evolving valence in the movie. Thus, an altered BOLD variability in these regions may reflect altered processing of emotional states during affective experience.

To investigate how this uncertainty around valence is computed by the young and older adults, we implemented a Bayesian learning model termed as Hierarchical Gaussian Filter model (HGF) (Figure. 5a). First, we collected the continuous valence ratings from two independent groups of young and older adults for the movie (Figure. 5b). Both the groups appeared to perceive the valence very similarly (*r* = 0.91, *p* < 0.001). Next, we deployed multiple 3-level HGF models to fit the averaged valence ratings of each group separately. This model has been extensively used in learning, decision-making, and affective dynamics^36,46,47^. In this model, the averaged valence time-course of each group is used as the input. With this input, the first layer, x_1_, models the probability of the valence at each time point. The next level, x_2_, estimates the underlying uncertainty associated with of the predictions on the valence. Finally, the third level x_3_, estimates the overall volatility or the metavolatility associated with the valence. This indicates the rate at which the belief on the uncertainty around valence changes. This can also be described as one’s belief in how stable are the movie contexts, given the valence dynamics. The belief updates are primarily governed by 2 parameters - ***ω*** and, which influence the estimated uncertainty (second-level) and volatility (third-level) trajectories about the valence respectively. Here, we wanted to dynamically model the averaged valence rating of the elderly, by lesioning these 2 free parameters - ***ω*** and ***ϑ***.

**Figure 5:**
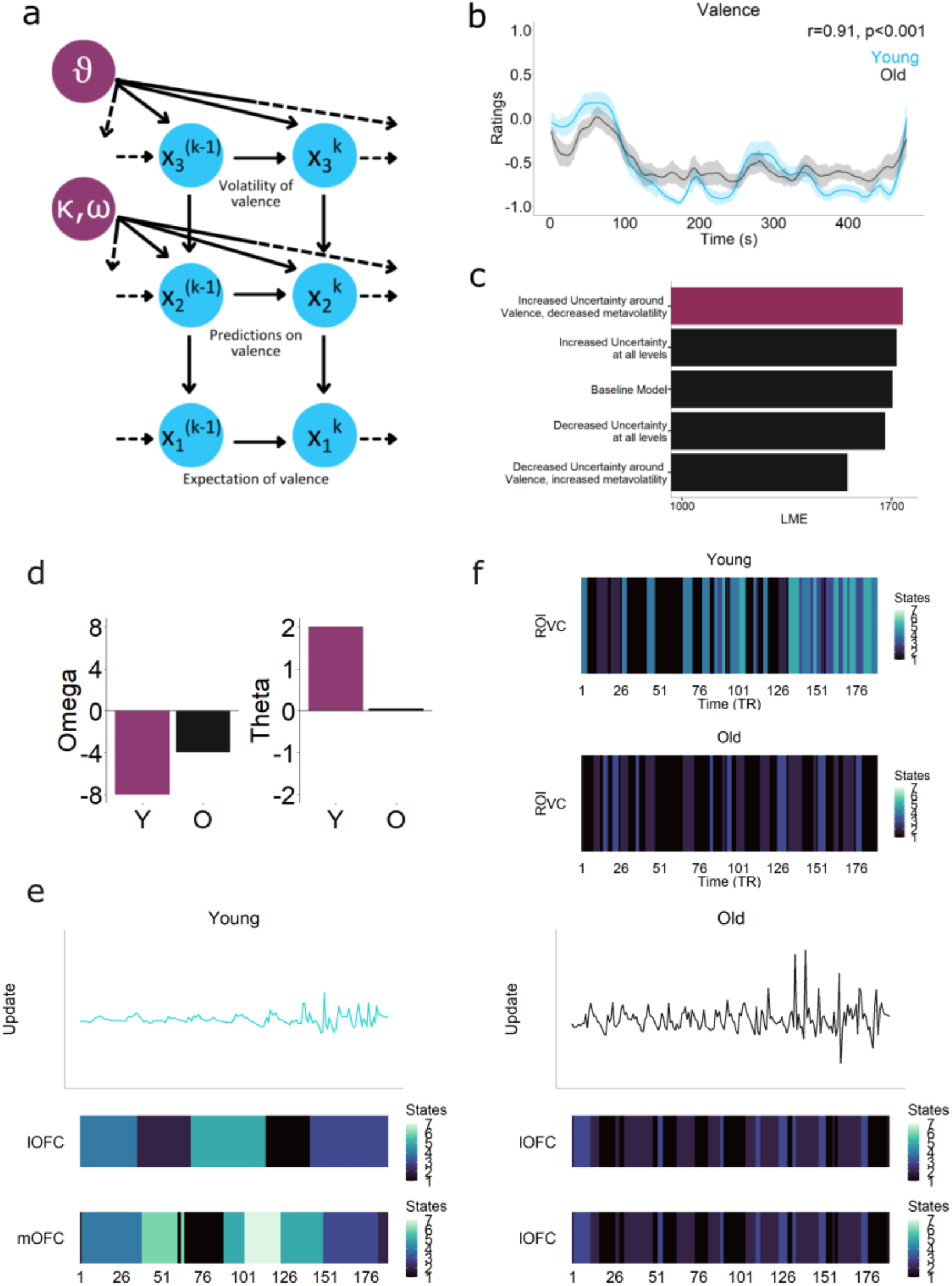
(a) Graphical illustration of the HGF. x_1_ represents beliefs about the observed valence, which in turn is affected by x_2_, representing the belief of how the valence changes. x_3_ estimates the metavolatility related to the valence timecourse, which in turn affects x_2_. The levels are coupled via a phasic, time-varying parameter kappa (κ). A tonic, time-invariant parameter omega (ω) influences the variance of the Gaussian walk at x_2_, while theta (***ϑ***) determines the variance of the volatility. (b) Continuous ratings of Valence during the Movie in young and older adults. Older adults exhibit more variable valence than young adults although they show a strong correlation (r=0.91, p<0.001). (c) We simulated multiple 3-level HGF models for older adults and selected the models based on the highest LME. These models were simulated by lesioning two free parameters - ω and ***ϑ*** affecting the second and the third level trajectories simultaneously. (d) Parameter comparisons from the final HGF model reveal an increased ω and reduced ***ϑ*** in the older adults. (e) Model prediction updates and neural state transitions of the lOFC and mOFC (from HMM). Younger adults show much more stable and synchronised latent state transitions of the OFC (qualitatively) which were replaced by much faster transitions in the older adults. (f) The latent state transitions of the Visual Cortex (VC) between the two groups, showed a similar transition pattern in both groups. This might reflect that the processing of sensory uncertainty (changing of visual features) is similar in both groups.

For this, we initially used the HGF model fitted to the young group (from our previous study)^20^ as the ‘baseline’ model. Following this, we simulated multiple models each with a specific lesioning in ***ω*** and ***ϑ*** parameters, which underscored various hypothesis of how their computations might differ (Figure. 5c). We then fit each of these models to the older cohort’s group-averaged valence time-course. We hypothesized that the model best-fitting the older adults’ task-specific variable (valence) would deviate from the ‘baseline’ parameters. This deviation would then reflect specific changes in the estimation of uncertainty around the valence, reflective of their observed differences in the neural variability of the OFC. A reduced ***ω*** in the older adults would suggest a decreased prediction uncertainty around the valence dynamics, thus slower updation to the changing valence. In contrast, an increased ***ω*** will signify increased uncertainty around the predictions of valence, thus faster updation to the changing valence. The meta-volatility parameter ***ϑ***, on the final level, determines the belief about the model’s beliefs in the movie volatility. Lower values denote higher beliefs in the stability of the movie context, and vice versa.

Model inversion was conducted by variational inference, and model adjudication was done by selecting the one with the highest log-model evidence (LME) (Figure. 5c) (Table 2). The best-fitting model showed an increased ***ω*** (Figure. 5d) suggesting an increased variance (uncertainty) around the valence in the elderly. Concurrently it also showed a decreased ***ϑ***, denoting an increased belief in the stability of the context of the movie. This suggests that the increased variability observed in the neural level (BOLD) might be reflective of higher uncertainty representations as a function of aging.

A more direct prediction of the model is that older adults would show more frequent prediction updates due to their increased ***ω*** (Figure. 5e right). That is, the belief update trajectories in the older adults showed more frequent updates compared to that of the young. Based on our previous findings that the OFC tracks uncertainty around valence during naturalistic movie viewing^20^, we hypothesized that older adults would show more frequent neural transitions in these regions compared to younger adults. We used Hidden Markov Models (HMMs) to obtain these neural transitions (see Methods for details). We found that both lOFC and mOFC showed more stable transitions in the young adults (Figure. 5e left). However, both these regions both showed more frequent latent state transitions in older adults (Figure. 5e right), linking the neural data with the model predictions.

We also compared the HMM state transitions of the visual cortex as a control ROI for the two groups (Figure. 5f). This was to rule out the effect of sensory or perceptual uncertainty processing. Both groups exhibited similar transitions with no distinct pattern, suggesting that the observed differences in orbitofrontal regions are due to top-down processes rather than perceptual differences.

In summary, an increased neural variability in older adults within OFC reflected representation of increased uncertainty during emotional processing. This also corresponded with faster latent state transitions within the older adults in contrast to more stable transitions within the OFC of the younger adults, aligning with frequent (Bayes-optimal) prediction updates.

### Spill-over effect of age-related emotional variability to multiple tasks

Next, we considered whether the heightened uncertainty during affective computations would result in more variable representations of emotions. It was important to rule out the possibility that the observed neural variability was specific to movie demands (plot complexity, working memory load etc). To this end, we analyzed data from the scanning cohort on a Facial Emotion Recognition task.

First, we examined a task in which participants reported the displayed emotion on a morphed image of a facial expression (FER). Participants chose from six emotion categories: Happy, Sad, Anger, Fear, Disgust, and Surprise. We compared the individual variability of each participant by computing the standard deviation (SD) of their accuracy over all 6 categories. Older adults had a significantly higher SD than young (mean-SD_Y_ = 11.08, mean-SD_O_ = 17.44, *t* = −5.17, *p* < 0.0001, Cohen’s *d* = 0.73) (Figure. 6a). Consistent with our earlier findings, between-group variability for each of the 6 emotions showed a similar pattern of increased idiosyncrasy in older adults (Happy: SD_Y_ = 2.68, SD_O_ = 5.99; Sad: SD_Y_ = 5.02, SD_O_ = 19.31; Anger: SD_Y_ = 19.52, SD_O_ = 27.4; Fear: SD_Y_ = 15.4, SD_O_ = 23.54; Disgust: SD_Y_ = 20.28, SD_O_ = 20.63; Surprise: SD_Y_ = 8.04, SD_O_ = 13.32) (Figure. 6b). These results indicate that emotional variability in older adults extends beyond the movie-watching context, emerging consistently across tasks and emotional categories – even under conditions with minimal narrative or memory demands.

**Figure 6:**
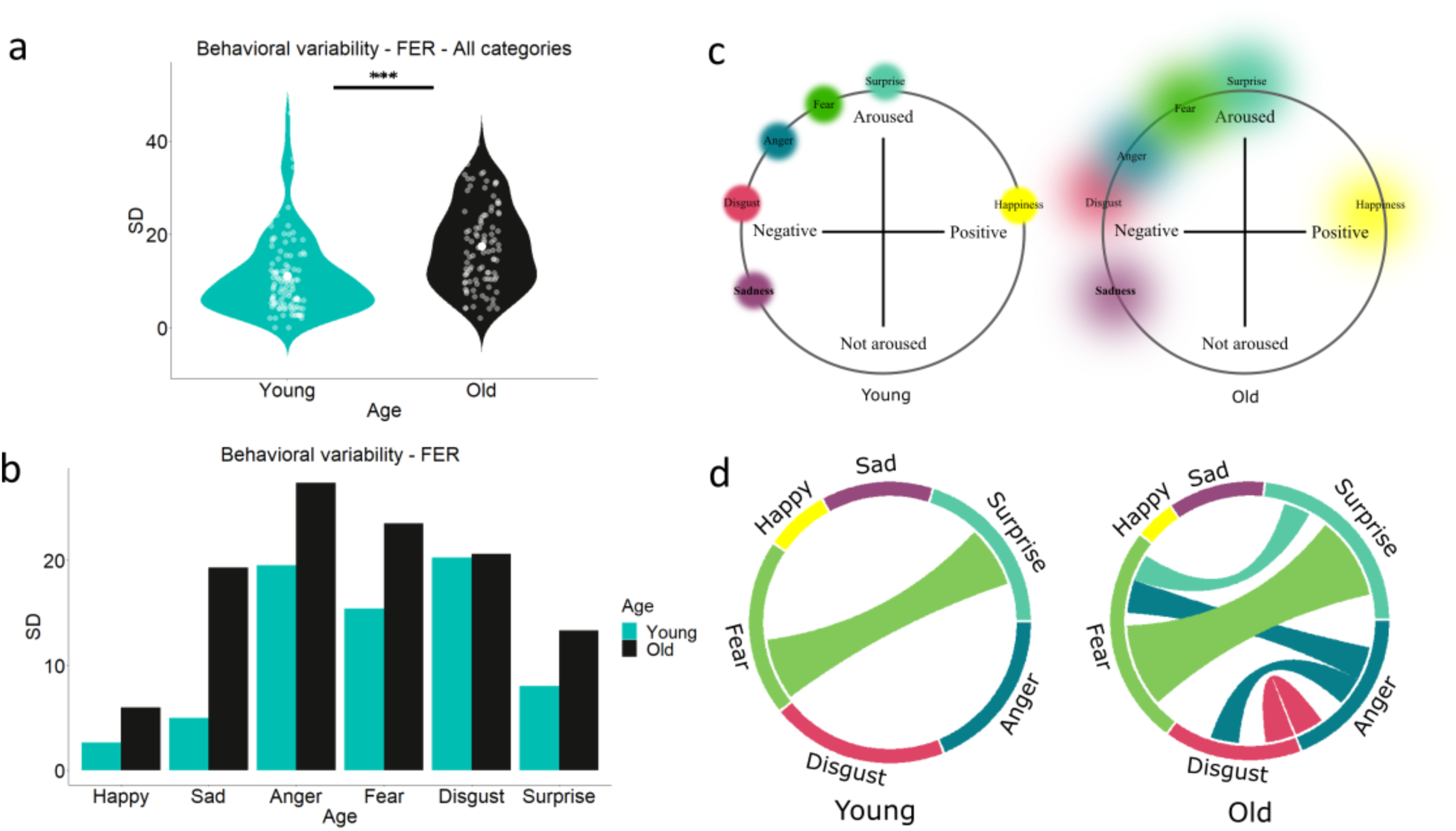
Generalizing age-related emotional variability to multiple tasks. (a) In the Facial Emotion Recognition (FER) task, individual variability, computed as the standard deviation (SD) of each participant’s accuracy over all 6 categories depicts significantly higher behavioral variability in older adults compared to young (mean-SD_Young_ = 11.08, mean-SD_Old_ = 17.44, *P*<0.0001). (b) Older adults display increased response variability compared to the younger individuals for each of the emotion categories. (c) We posited that increased variability (represented as coloured circles around emotion categories) around the valence leads to the misrepresentation of emotions in older adults. (d) Chord diagram showing a “spill-over” effect of adjacent categories in older adults.

Considering the classic circumplex model of emotions (Figure. 6c) an increased variability around an emotion not only would compromise accurate perception but also result in a systematic ‘spill-over’ of adjacent emotions (but not distant ones) into each other, contributing to a highly variable emotional profile that may have signatures in cortical regions involved. So, we explored whether this variability underlies any specific pattern of responses across the categories. For this we computed a co-occurrence matrix of the true emotion (of a face) and the responded emotion by the participants, for each group. For example, if, out of 100 instances of Fear trials, 80 accurate Fear responses were paired with 20 inaccurate Surprise responses, we could infer that Surprise had a ‘spill-over’ effect into the Fear category. For visualization, we collapsed it across participants for all six emotion categories for each group (values adjusted based on a predetermined threshold). Figure. 6d shows that older individuals exhibited substantial bidirectional overlap between surprise and fear indicating significant overlapping representation of these two emotions. Similar patterns are observed between anger and disgust, as well as between anger and fear. Notably, these emotions are adjacently placed in the circumplex model. If the observed overlap were solely a result of their variability in representing faces, one would expect equal overlap among all emotions, which is not the case. Likewise, this phenomenon cannot be purely attributed to the variability in representing negative emotions in general, as it fails to explain the specificity observed between adjacent emotions (e.g. no overlap between anger and sadness, which are positioned farther apart than disgust or fear). The young adults showed significant overlap only between fear and surprise.

Taken together, these findings indicate that older adults exhibit increased variability in emotion recognition across all categories, with specific patterns of overlap between certain adjacent emotions.

## Discussion

The key advance of our work is to comprehensively demonstrate that neural volatility captured via temporally organized regional BOLD signal variability during naturalistic tasks is a non-trivial, physiologically distinct marker of age-associated changes in affective computations. This was not shown before by any previous studies which has investigated how volatility facilitates value updating in the frontal cortex, adding to the novelty of these findings and broader implications of volatility during emotion processing under dynamic contexts and tasks. Importantly, the changes to cortical neural variability in older adults revealed distinct patterns during naturalistic experience, which were neither captured by the mean neural activity nor the resting-state variability from the same individual. This is in line with theoretical frameworks that posit task-variable computations require navigating different regimes flexibly, which may be compromised in aging^15^. We found during naturalistic emotion processing; older adults exhibit an increased neural volatility uniquely localized in the mOFC and lOFC compared to other prefrontal subregions. Crucially, the representation of latent state trajectories relevant to the observed volatility in these regions closely resembled the dynamic affective experiences of the two groups during the movie-watching task. Increased neural volatility in the OFC was also associated with reduced positivity bias (widely reported in the aging literature), suggesting an adaptive link between neural volatility and individual’s affective profiles. Moreover, to understand how the affective dynamics optimally depend on the level of uncertainty or volatility, a computational modelling framework employing a Bayesian learning model further revealed that this volatility in older adults was associated with an increased uncertainty around valence computations. This increase in volatility predicted more frequent updates, which was observed in OFC neural state transitions in the old compared with the younger group. Complementary evidence from a separate facial emotion recognition task on the same participants demonstrated that emotional variability in older adults extends beyond naturalistic movie viewing, emerging consistently across tasks and affective categories, even in contexts with minimal narrative or memory demands. These results advance our understanding of how neural volatility might carry unique information in aging, particularly with regard to their idiosyncratic affective experiences.

Neural variability is usually studied on cognitively demanding tasks (e.g. working memory) or resting-state variability^9,10,14,15,17^. These studies provide a wealth of results in terms of its various neurobiological properties and alterations in healthy aging^48^. While some studies link increased neural volatility to deteriorating performance in tasks like working memory^11^, others demonstrate the opposite trend. The pattern of change also varies across different brain regions. Unlike specific task-based studies, naturalistic paradigms require coordinated dynamics across multiple forms of higher-order cognition, making them more ecologically similar to real-life experiences^49,50^. Leveraging Bayesian hierarchical modelling allowed us to incorporate multiple, related sources of information within the data. Specifically, this ensured that we captured both individual, group-level and condition-specific effects on neural volatility. In doing so, we found a significant increase in neural volatility in older adults compared to young. Critically the evidence for this increase was much smaller in intrinsic, resting-state BOLD variability. This comparison provides strong evidence that neural variability modulated by naturalistic experiences differs from spontaneous, resting-state activity. In contrast, the mean neural responses (BOLD mean) showed no differences among the young and old. This strongly suggests that the mean and SD of the physiological BOLD signal carry fundamentally different sources of information in the context of emotion processing in aging^48^.

Naturalistic stimuli evoke rich and dynamic patterns of neural activity, integrating multiple cognitive and affective processes. In this study, we focused on continuous valence as a key dimension of emotional experience. In addition, we focused on the OFC because of its critical role in affective processing^20,21,37,38^. Studies have shown that its lateral subregions (lOFC) underpin distinct functional roles than its medial counterparts (mOFC)^51,52^. Here, we observed a consistent increase in neural variability in both lOFC and mOFC in older adults compared to the younger individuals. Importantly, these differences in volatility could not be attributed to the differences in perceptual processing in the visual cortex. No differences were seen in volatility in prefrontal regions involved in social processing (mPFC) and attentional deployment (dlPFC), both of which are key in movie processing among the groups (Figure. 2c, d). Moreover, the mean neural activity showed no differences among the groups in both regions. These results suggest that OFC neural volatility during movie viewing uniquely captures age-related alterations in affective processing.

A unique advantage of naturalistic stimuli is its deep ties with affective experience, embedded in cortical latent dynamics. We assumed changes to OFC volatility would be reflective of underlying changes to affective computations specific to the movie. This change should be then reflected in the underlying latent representations over time. Using T-PHATE, we projected neural representations of the OFC regions onto a low-dimensional space, uncovering these latent trajectories across the two groups. The stark differences observed in the lOFC and mOFC were highly similar to their respective arousal time-courses. That is, segments of the movie where affective computations were more demanding, reflected in increased Arousal, had distinctly different trajectories in the OFC. Specifically, a sequentially preserved representation observed in the young group mirrored their respective stable arousal dynamics. In contrast, the distorted patterns observed in older adults’ lOFC and mOFC paralleled their lower and more variable arousal.

Do these findings reflect responses specific to movie viewing, or do they generalize to broader affective computations in aging? If neural volatility is an adaptive response to aging, where older participants are compensating for computational deficits, then it might be related to affective behavior as a whole. To test this, we examined behavior in an independent emotional reactivity task, where same participants passively viewed short emotional clips and rated their felt affect. Older adults are typically characterized by a bias toward positive stimuli ^41^, though this effect is variable across contexts^43^. In this study we found that movie-evoked neural volatility in both the lOFC and mOFC was significantly related to this positivity bias where individuals with greater OFC volatility exhibited a reduced bias toward positive affect. This relationship was specific to the OFC and was not observed in other prefrontal regions implicated in processing social or semantic content in the movie. These findings suggest that OFC volatility reflects broader alterations in affective computations with age and may serve as a neural marker of individual differences in emotional bias.

Theoretical models posit that neural volatility represents optimal task computations and uncertainty representations^4,12,13^. Optimally representing task-uncertainty comes under the purview of Bayesian frameworks of cognition^53^. To explore the effect of increased neural volatility at the computational level during naturalistic affective experience, we used a hierarchical Bayesian learner ^36^ which was given the valence responses in the movie. In our previous study, a normative variant of this model showed how uncertainty computations arise from valence in the same younger individuals^20^. Using this as our baseline, we adjusted two key parameters underlying uncertainty or agent volatility computations in different ways. This was to test several competing accounts of suboptimal computations happening in aging^46,54,55^. This allowed us to use the same model architecture to assess deviations in older adults compared to young. The winning variant showed considerably higher variance around predicting valence for the older adults. That is, older adults had a heightened volatility around predicting how the valence changes in the movie. To link the neural findings with the model output, we compared prediction updates in the model to that of the latent state transitions in the OFC. Neural state changes during naturalistic stimuli are closely linked to the movie experience ^20,47^, although several factors might underlie these changes within a region. Here, we found qualitative similarities between OFC transitions and prediction updates to valence in the young. In contrast, the older adults showed more frequent neural transitions. This suggests that neural transitions here may reflect an increased estimation of uncertainty around valence in these regions. Overall, increased uncertainty around the valence coupled with increased neural volatility in OFC suggests that older adults may experience less stable or more variable emotional representations.

Accurate emotion recognition, such as identifying facial expressions, likely depends on stable internal representations of distinct emotional categories. Aging may disrupt these representations, in part due to increased uncertainty, leading to more frequent misclassifications. Heightened uncertainty around one emotion category (depending on the internal representations), then might show forms of ‘spill-over’ to adjacent emotion categories more than distantly placed ones (Figure. 6c,d). Our results from the facial emotion recognition task support this interpretation, revealing a highly variable and idiosyncratic emotional profile in older adults. Such spill-over effects suggest a systematic distortion in the 2D emotional space, potentially arising from altered neural encoding in cortical regions involved in affective processing. Of course, other sources such as allocation of attention to only some facial features potentially might be affecting this partly or jointly. Future studies could implement eliciting specific emotions without the medium of faces to confirm it is not due to attentional allotment to facial features alone.

Cognitively demanding tasks, such as working memory, require representing and navigating a higher dynamic range of states to perform optimally. Affective tasks, in contrast, are more inferential^56^. As a result, there is an inherent uncertainty in one’s interpretations of emotional experiences. Here, representing a wider gamut might be misleading (at best) or paralyzing (at worst). Indeed, misestimation of uncertainty, a known factor in psychiatric disorders ^28,57^, may similarly underlie the idiosyncratic affective experiences in aging. In affective tasks, the lOFC helps predict how emotional states will change over time, integrating information about uncertainty to guide decision-making^22,26^. The higher volatility observed here in the older cohorts seems to be consistent with our model results and reward learning studies. Taken together, this suggests that a failure to compute the uncertainty around the valence by the OFC might underlie the consistently idiosyncratic affective experiences seen in aging.

There are inherent limitations to interpreting BOLD signal volatility in any task. For example, vascular and non-cognitive processes may play a significant role^48^. We took steps to mitigate these influences by regressing motion as well as the CSF and white matter signals. Another potential critique is the use of two separate behavioral and neuroimaging groups in this study. Prior works^33,58^ have obtained ratings and neural time series from two cohorts and used group-averaged estimates of the former to investigate the latter, similar to our approach. Moreover, we generalized the results of variability in emotions in the scanning cohort to two separate tasks requiring the processing of valence, as well as link their behavior to neural variability.

Overall, we put forward a strong case for the physiological BOLD signal variability as a proxy of neural volatility, as a readout of regional computations, separate from resting-state and neural mean activity. The changes to this signal in healthy aging, captured in a naturalistic setting, offered a window to their idiosyncratic experience. More nuanced insights on how various forms of uncertainty can affect the volatility remain open questions, however, we hope this study has opened avenues and offered a novel framework to address such questions by future studies.

## Acknowledgments

NBRC Core funds supported this study. DR was supported by SERB Core Research Grant (CRG) S/SERB/DPR/20230033 extramural grant from the Department of Science and Technology, Ministry of Science and Technology, Govt. India. DR and AB acknowledge the generous support of the NBRC Flagship program BT/MIDI/NBRC/Flagship/Program/2019: Comparative mapping of common mental disorders (CMD) over the lifespan. Further, we would sincerely like to thank Rindish Krishna for his technical help in developing the software for collecting behavioral responses. We would also thank Rishi Dey Chowdhury for his involvement in implementing the Temporal PHATE analysis in this study.

