## Extended Data for "Neural volatility in orbitofrontal cortex underlies affective processing during healthy cognitive aging"

### Supplementary results

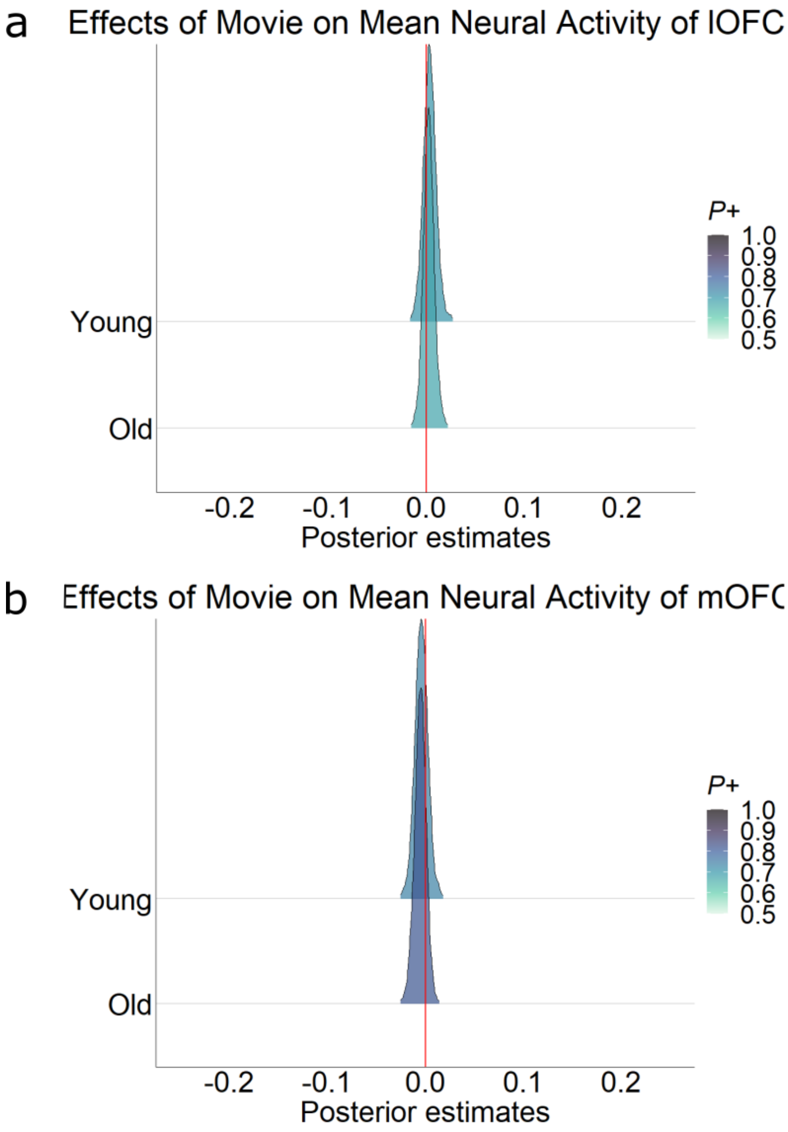


**Figure 1-1:** Mean neural activity of both lOFC and mOFC shows no significant difference between the two groups for the movie-watching task. ROI-level effect of the movie on the $\mu_{BOLD}$ (averaged BOLD signal) across the two groups. Posterior density estimates of (a) lOFC and (b) mOFC shows no significant difference between the two groups during movie.

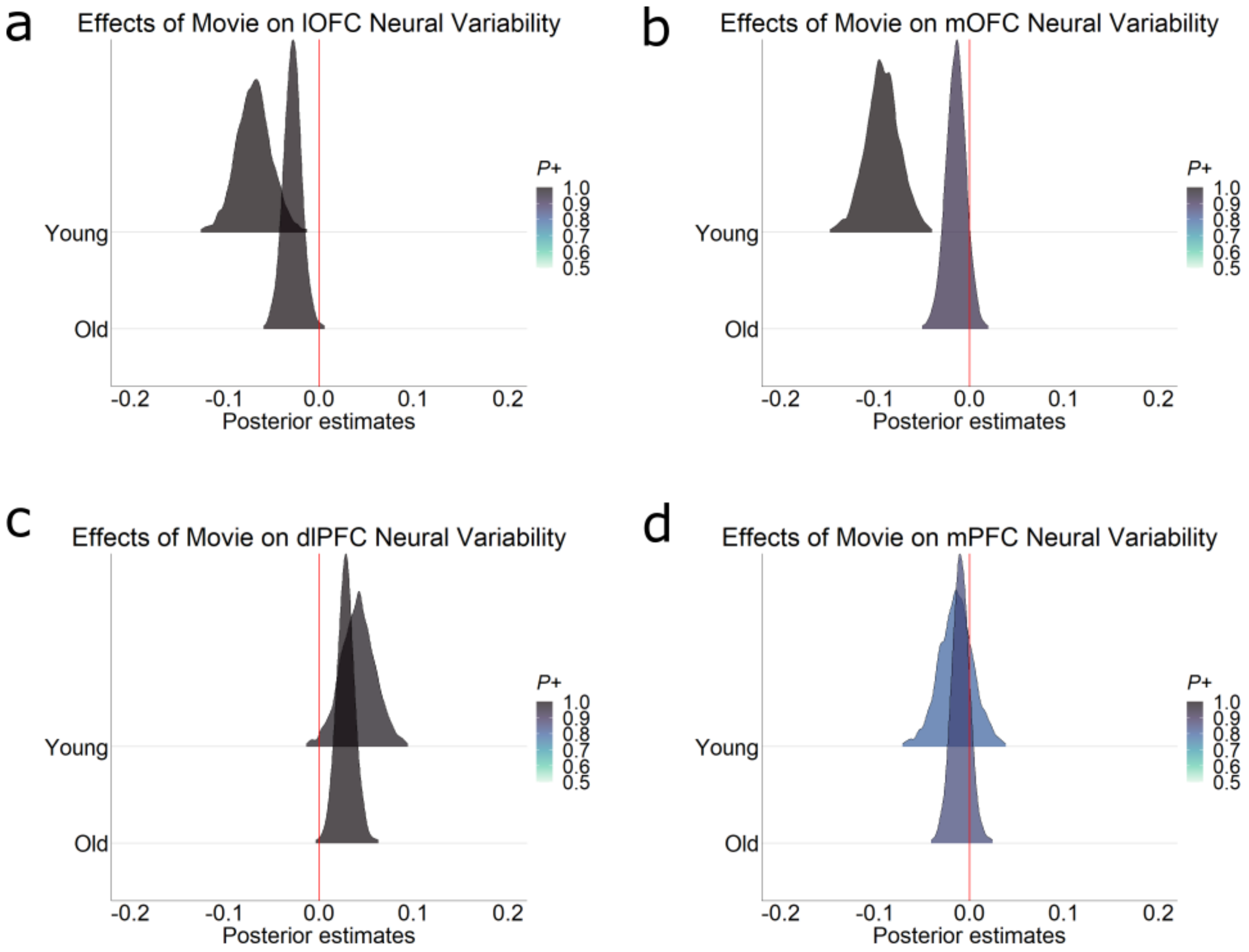


**Figure 2-1:** Neural Variability shows functionally distinct trend in the Orbitofrontal cortex in aging (with ICA-AROMA). ROI-level effect of the movie on the $SD_{BOLD}$ across the two groups. (a) Posterior density estimates of lOFC shows significantly more evidence in old than young (Young: $\beta_{mean} = -0.07$, $P = 0.9$; Old: $\beta_{mean} = -0.03$, $P = 0.9$). (b) The posterior density estimates of mOFC showed similar pattern of increased variability in old compared to younger adults (Young: $\beta_{mean} = -0.09$, $P = 1$; Old: $\beta_{mean} = -0.01$, $P = 0.9$). No significant differences were observed in the neural variability between the two groups for (c) the dlPFC (Young: $\beta_{mean} = 0.04$, $P = 0.9$; Old: $\beta_{mean} = 0.03$, $P = 0.9$) and (d) mPFC (Young: $\beta_{mean} = -0.01$, $P = 0.7$; Old: $\beta_{mean} = -0.01$, $P = 0.8$).

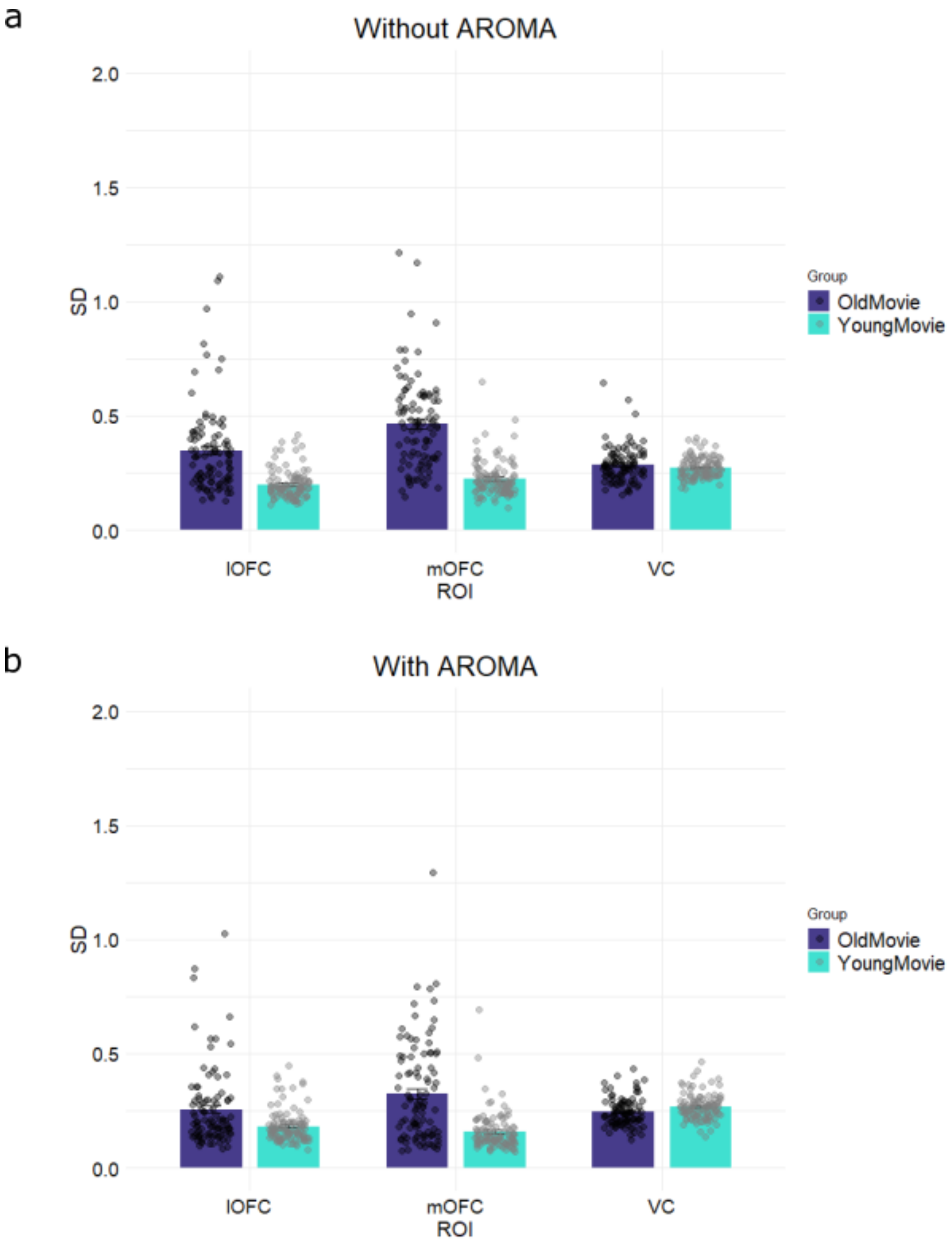


**Figure 2-2:** Comparison of the $SD_{BOLD}$ for two separate preprocessing pipelines with and without ICA-AROMA. The lOFC and mOFC in both cases show significantly higher $SD_{BOLD}$ in older

adults compared to young. In contrast the VC shows insignificant differences between the two groups.

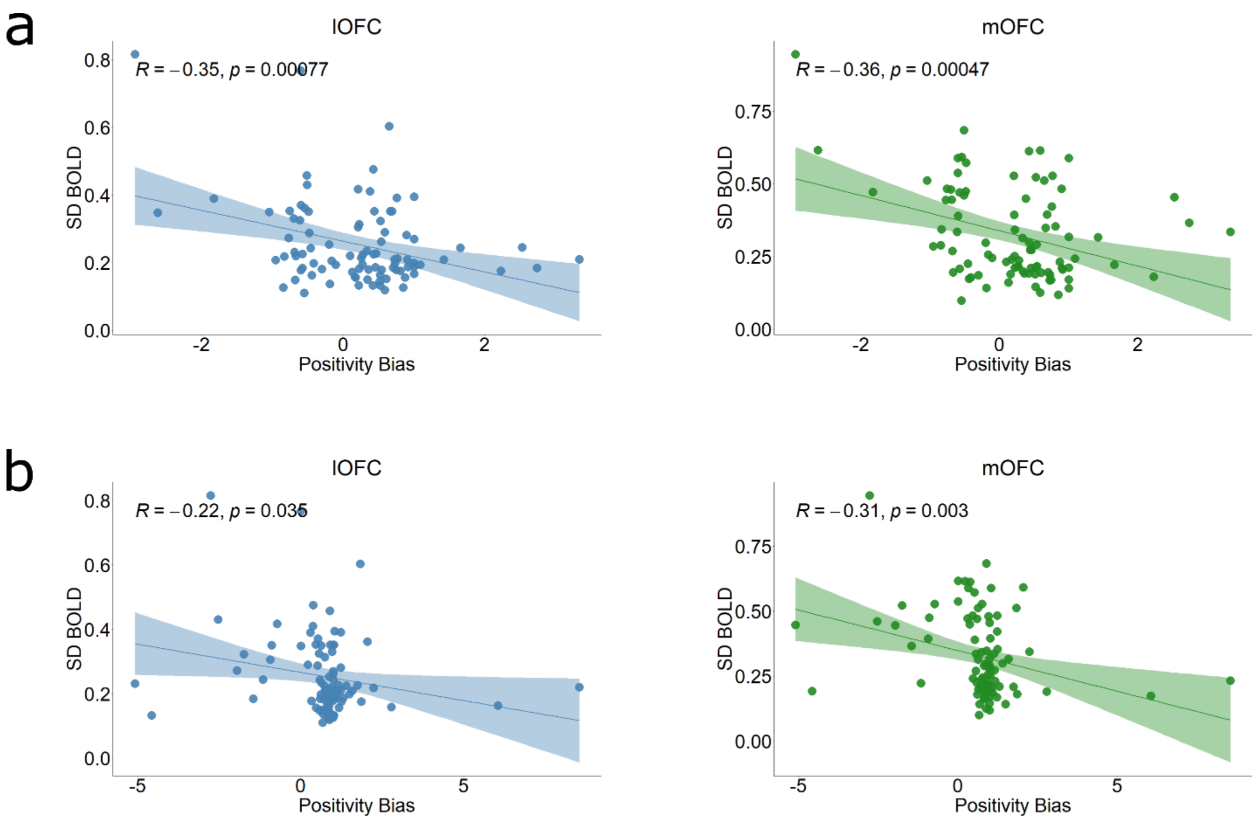


**Figure 4-1:** (a) Increased neural variability in the orbitofrontal cortex is associated with reduced positivity bias in aging for Neutral clips and (b) Negative clips. The positivity bias is computed as the ration of the positive to negative ratings for each Condition (Neutral and Negative).
